# Wnt/β-catenin signaling restrains developmental beige adipocyte thermogenesis and its inhibition imprints long-term energy expenditure

**DOI:** 10.64898/2026.03.23.713637

**Authors:** Ganran Xu, Tian Chen, Qiuwen Zhang, Bohuai Zhou, Jingfan Xue, Jingang Xiao, Ding Bai, Yiping Chen, Weidong Tian, Zhi Liu

## Abstract

Beige adipocytes that emerge during the peri-weaning period support sympathetic nervous system (SNS)-independent thermogenesis, yet the mechanisms governing this spontaneous beiging remain unclear. Here, by integrating transcriptomic profiling with adipocyte-targeted *Ctnnb1* deletion in mice, we identify the canonical Wnt signaling as an endogenous brake on developmental beige thermogenesis. Peri-weaning inguinal fat from adipocyte *Ctnnb1* knockout mice exhibits enhanced beige adipocyte biogenesis, with increased thermogenesis-related gene expression and mitochondrial oxidative capacity, which programs durable activation of adaptive thermogenesis and augmented whole-body energy expenditure into adulthood. Mechanistically, suppression of Wnt/β-catenin signaling induces a non-canonical Wnt5a-Ca^2^⁺-AMPK axis that promotes triglyceride lipolysis and subsequent PPAR-driven fatty acid oxidation, thereby fueling mitochondrial respiration. Genetic or pharmacological disruption of this axis blunts thermogenic responses induced by β-catenin inhibition in both murine and human subcutaneous adipocytes, indicating that Wnt5a-Ca^2^⁺-AMPK axis is required for the cell-autonomous activation of beige fat. These results reveal Wnt/β-catenin signaling as a developmental constraint on beige adipocyte formation and suggest an SNS-independent route to sustainably raise energy expenditure and improve metabolic health.

## Introduction

Adipose tissue plays a pivotal role in energy balance and systemic metabolic regulation, serving not only as the principal site for energy storage but also as a key contributor to thermogenesis^1,2^. Thermogenic adipocytes are specialized fat cells that dissipate chemical energy as heat and are consistently linked to improved glucose homeostasis, enhanced lipid clearance, and protection against obesity-related metabolic disorders^3,4^. Two types of thermogenic adipocytes, known as brown and beige adipocytes, have been characterized by the expression of uncoupling protein 1 (UCP1), a key thermogenic marker^5,6^. Brown adipocytes, residing in brown adipose tissue (BAT), are largely constitutive throughout life, while beige adipocytes can be recruited within subcutaneous white adipose tissue (sWAT) in response to external cues such as environmental cold acclimation^5,7–9^. In adult mammals, beige adipocyte recruitment is predominantly driven by sympathetic nervous system (SNS)-mediated β-adrenergic receptor (β-AR) signaling, with intracellular regulators such as AMP-activated protein kinase (AMPK) and peroxisome proliferator-activated receptors (PPARs) modulating the thermogenic program^2,3,10–16^. However, this reliance on sustained SNS input limits the feasibility and durability of beige adipocyte activation as a therapeutic strategy. Moreover, pharmacological β3-AR agonists, although capable of stimulating beige fat, have been associated with increased cardiovascular risk^11,17–19^. Thus, alternative mechanisms that promote beige adipocyte formation and function independent of SNS activation are of considerable interest.

Emerging evidence indicates that, in contrast to the adult situation, beige adipocyte biogenesis during development can occur independently of environmental temperature and sympathetic stimulation^20–26^. In mice, sWAT undergoes spontaneous beige adipogenesis during the peri-weaning period without cold exposure, and this process persists even under thermoneutral conditions or after sympathetic denervation^24^. The Ucp1^+^ beige adipocytes generated during this developmental window subsequently regress into a white-like state, referred to as dormant or latent beige adipocytes, which remain highly plastic and can be rapidly reactivated by later environmental challenges^20^. These dormant cells thereby constitute a reservoir of thermogenic potential to support adaptive thermogenesis and confer long-term metabolic benefits into adulthood^20,24,27–29^. Furthermore, experimental evidence suggests that increasing the pool of dormant beige adipocytes enhances systemic metabolism even in the absence of thermogenic activation^24,27,28^. Despite their physiological significance, the molecular signals governing the spontaneous emergence of peri-weaning beige adipocytes remain poorly understood.

In this study, we took advantage of the peri-weaning window of spontaneous beige adipocyte emergence to probe the intrinsic signals that regulate developmental beige thermogenesis. On this basis, we identify the canonical Wnt signaling as a previously underappreciated endogenous brake on this process, as its inhibition selectively enhances beige adipocyte activity and mitochondrial oxidative capacity in sWAT, but not in BAT or epididymal WAT (eWAT). Furthermore, attenuating Wnt/β-catenin signaling does not overtly impair adipocyte differentiation or endocrine function but instead biases lipid handling towards increased triglyceride lipolysis to fuel mitochondrial thermogenesis. Importantly, adipocyte *Ctnnb1* ablation establishes a higher thermogenic tone in subcutaneous depots and leads to sustained increases in whole-body energy expenditure into adulthood. Mechanistic analyses indicate that Wnt/β-catenin signaling exerts its restraining effect, at least in part, by suppressing a non-canonical Wnt5a-Ca^2^⁺-AMPK axis. Genetic and pharmacological disruption of this axis abolishes the thermogenic response to β-catenin inhibition in both murine and human subcutaneous adipocytes. Notably, Wnt5a overexpression alone fails to activate AMPK or induce thermogenesis when Wnt/β-catenin signaling remains intact, indicating that release of the canonical Wnt brake is required for this non-canonical axis to drive beige adipocyte activation. Together, these findings define Wnt/β-catenin signaling as a developmental constraint on spontaneous beiging in sWAT and point to a potential SNS-independent route to improve metabolic health.

## Results

### Peri-weaning iWAT adipocyte precursors display reduced Wnt/β-catenin signaling and higher thermogenic capacity upon differentiation

Developmental beige adipocytes, which become thermogenic inactive in adulthood, have been reported to transiently and spontaneously arise in sWAT of young mice around the peri-weaning period under ambient-temperature conditions^21–26^. To delineate the temporal pattern of this phenomenon, we profiled thermogenic features of the inguinal white adipose tissue (iWAT), a major subcutaneous depot in mice, at room temperature across age. Analysis of wild-type male mice at 3, 4, 5, and 8 weeks of age revealed a pronounced peri-weaning peak in thermogenesis-related markers around 4 weeks, followed by a progressive decline (Fig. 1a,b, Supplementary Fig. 1a). We next asked whether this developmental beiging is driven primarily by age-dependent systemic cues or by cell-autonomous properties of adipocyte precursors. To this end, we compared the thermogenetic capacity of adipocytes differentiated in vitro from the stromal vascular fraction (SVF) cells isolated from peri-weaning (4-week) and adult (8-week) iWAT. Under the standard white fat pro-adipogenic induction, adipocytes derived from 4-week mice displayed smaller, multilocular lipid droplets and significantly higher levels of thermogenic genes and proteins than those from 8-week mice (Fig. 1c–e, Supplementary Fig. 1b). These observations suggest that peri-weaning adipocyte precursors are intrinsically biased towards a more thermogenic phenotype than their adult counterparts under induction.

**Fig. 1.**
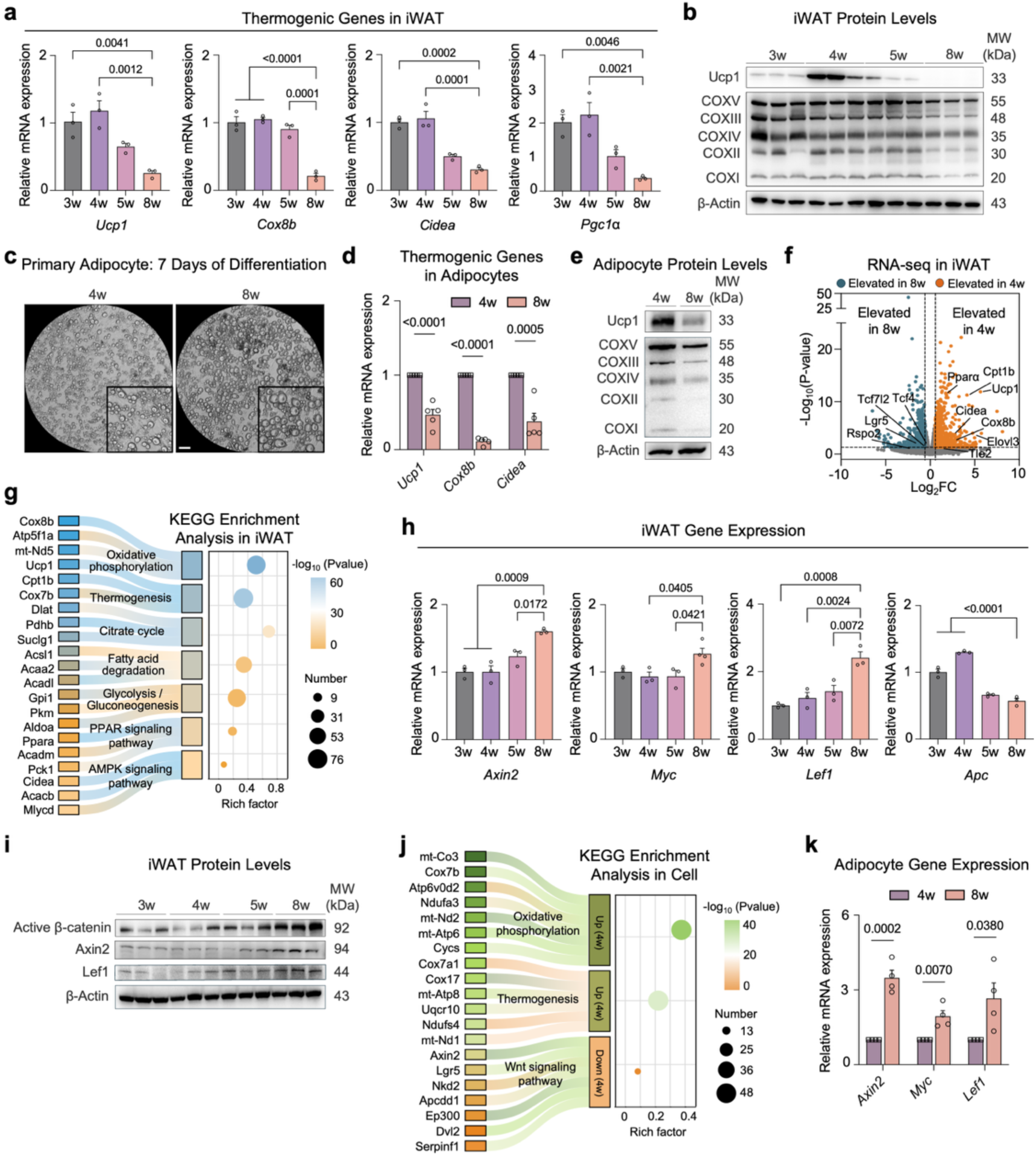
Peri-weaning iWAT adipocyte precursors display reduced Wnt/β-catenin signaling and higher thermogenic capacity upon differentiation. **a** mRNA expression of thermogenic genes in iWAT of wild-type male mice at 3, 4, 5, and 8 weeks of age (*n* = 3). **b** Immunoblotting for Ucp1 and OXPHOS in (a) (*n* = 3). c–e SVFs isolated from iWAT of wild-type mice at 4 and 8weeks of age were differentiated into adipocytes after 7 days. **c** Representative images of differentiated adipocytes. Scale bar, 200 μm. **d** mRNA expression of thermogenic genes in (c) (*n* = 5 biological replicates). **e** Immunoblotting for Ucp1 and OXPHOS in (c) (*n* = 3 biological replicates). **f,g** Data was processed using DESeq2 software for normalization and differential analysis, with a threshold of |FC| > 1.5 and adjusted P-value < 0.05. RNA-seq was performed in iWAT from wild-type mice at 4 and 8weeks of age (*n* = 3). **f** Volcano plot representing genes differentially expressed. **g** KEGG enrichment analysis of genes upregulated in 4-week mice relative to 8-week mice. **h,i** mRNA expression (**h**) and Immunoblotting (**i**) in iWAT of wild-type mice at 4 and 8weeks of age (*n* = 3). **j** RNA-seq was performed in (**c**), with a threshold of |FC| > 1.5 and adjusted P-value < 0.05 (*n* = 3). KEGG enrichment analysis of genes differentially expressed (*n* = 3). **k** mRNA expression of downstream genes of Wnt/β-catenin signaling pathway in (**c**) (*n* = 4 biological replicates). The levels of mRNA expression are normalized to that of *36B4*. Data are the mean ± s.e.m. Statistical analyses used were one-way ANOVA with Tukey’s correction for multiple comparisons or unpaired two-tailed *t*-tests.

To explore the molecular basis underlying this developmental difference in thermogenic potential, we performed bulk RNA-sequencing (RNA-seq) on iWAT from 4-week and 8-week mice. Differential gene expression analysis visualized as volcano plots revealed robust upregulation of genes associated with thermogenesis and oxidative metabolism, such as *Ucp1* and *Cpt1b*, in 4-week iWAT (Fig. 1f). Consistently, the Kyoto Encyclopedia of Genes and Genomes (KEGG) pathway enrichment analysis showed that pathways related to oxidative phosphorylation, thermogenesis, fatty acid metabolism were preferentially enriched in 4-week iWAT (Fig. 1g), supporting a more active thermogenic and energy-demanding state in the peri-weaning iWAT.

Notably, we also observed reduced expression of several genes with respect to Wnt/β-catenin signaling transcriptional activity, including *Tcf4*, *Lef1*, *Lgr5*, *Rspo2*, and *Tle2*, in 4-week compared with 8-week iWAT (Supplementary Fig. 1c), implying lower levels of Wnt/β-catenin signaling in the peri-weaning depot^30^. To validate this observation, we analyzed murine iWAT from 3 to 8 weeks of age, which showed consistently lower expression of the canonical Wnt pathway components together with reduced β-catenin protein abundance during the peri-weaning period compared with adulthood (Fig. 1h,i, Supplementary Fig. 1d). To further resolve whether Wnt pathway activity is spatially heterogeneous within iWAT, we profiled Wnt/β-catenin signaling activities across anterior, middle and posterior subregions of wild-type depots using *Axin2*, a β-catenin/TCF target gene, as a readout. In 4-week iWAT, *Axin2* expression was significantly higher in the posterior region than in the anterior and middle regions (Supplementary Fig. 1e), aligning with the observation that the posterior portion exhibits much weaker spontaneous beiging at this age (Supplementary Fig. 1f). Moreover, by 8 weeks of age, *Axin2* expression in the anterior and middle regions strongly increased and approached posterior levels under ambient conditions (Supplementary Fig. 1g), consistent with a more uniformly elevated Wnt/β-catenin tone at an age when spontaneous beiging in iWAT is markedly attenuated.

To minimize potential confounding from changes in tissue composition and non-adipocyte populations, we performed RNA-seq on adipocytes differentiated in vitro from SVFs isolated at 4 and 8 weeks. Consistent with the tissue-level data, 4-week SVF-derived adipocytes exhibited higher expression of thermogenic and oxidative genes alongside reduced Wnt pathway activity (Fig. 1j). This reduction in Wnt/β-catenin signaling was further corroborated by decreased expression of β-catenin/TCF target genes, including *Axin2*, *Myc*, and *Lef1*, in SVF-derived adipocytes in vitro (Fig. 1k). Collectively, these results indicate that the robust peri-weaning beiging phenotype is accompanied by reduced Wnt/β-catenin signaling activity in subcutaneous fat.

### Wnt/β-catenin signaling modulates developmental beiging

Next, we asked whether reducing Wnt/β-catenin signaling specifically in adipocytes would be sufficient to modulate beiging activity in vivo. For this purpose, we generated adipocyte-specific *Ctnnb1*-knockout mice upon compounding *Adipoq*^Cre^ allele with floxed *Ctnnb1* alleles (*Adipoq*^Cre^;*Ctnnb1*^flox/flox^), using *Adipoq*^Cre^;*Ctnnb1*^flox/+^ littermates as controls (Supplementary Fig. 2a). To evaluate recombination specificity, we separated adipose tissues from *Adipoq*^Cre^;*Ctnnb1*^flox/flox^ and control mice into SVF and mature adipocyte fractions (Supplementary Fig. 2b). We found that, across iWAT, BAT, and eWAT, *Ctnnb1* expression was efficiently suppressed in mature adipocytes but remained unchanged in SVF cells and in other tissues (Supplementary Fig. 2c,d). Consistently, β-catenin protein levels were markedly decreased in iWAT from 4-week-old *Adipoq*^Cre^;*Ctnnb1*^flox/flox^ mice, confirming effective *Ctnnb1* deletion in adipocytes (Supplementary Fig. 2e).

We then investigated the thermogenic phenotype of *Adipoq*^Cre^;*Ctnnb1*^flox/flox^ and control mice at 4 weeks of age. *Adipoq*^Cre^;*Ctnnb1*^flox/flox^ mice displayed significantly lower body weight than littermate controls (Fig. 2a), and their iWAT depots appeared smaller and more reddish (Fig. 2b). Histological analysis of iWAT revealed that adipocytes from *Adipoq*^Cre^;*Ctnnb1*^flox/flox^ mice, particularly in the posterior region relative to controls, exhibited a beige-like morphology with reduced lipid droplet size and a characteristic multilocular architecture (Fig. 2c, Supplementary Fig. 3a). At the molecular level, *Adipoq*^Cre^;*Ctnnb1*^flox/flox^ mice exhibited strongly enhanced iWAT beiging compared with control littermates, as evidenced by increased expression of thermogenic genes and elevated UCP1 and OXPHOS protein abundance (Fig. 2d–f, Supplementary Fig. 3b,c). In parallel, β-catenin deficiency significantly increased the respiratory capacity of iWAT, including both basal and maximal oxygen consumption (Fig. 2g, Supplementary Fig. 3d). These data indicate that loss of β-catenin in adipocytes during the developmental period augments beige adipocyte activation and enhances oxidative metabolism in iWAT.

**Figure. 2.**
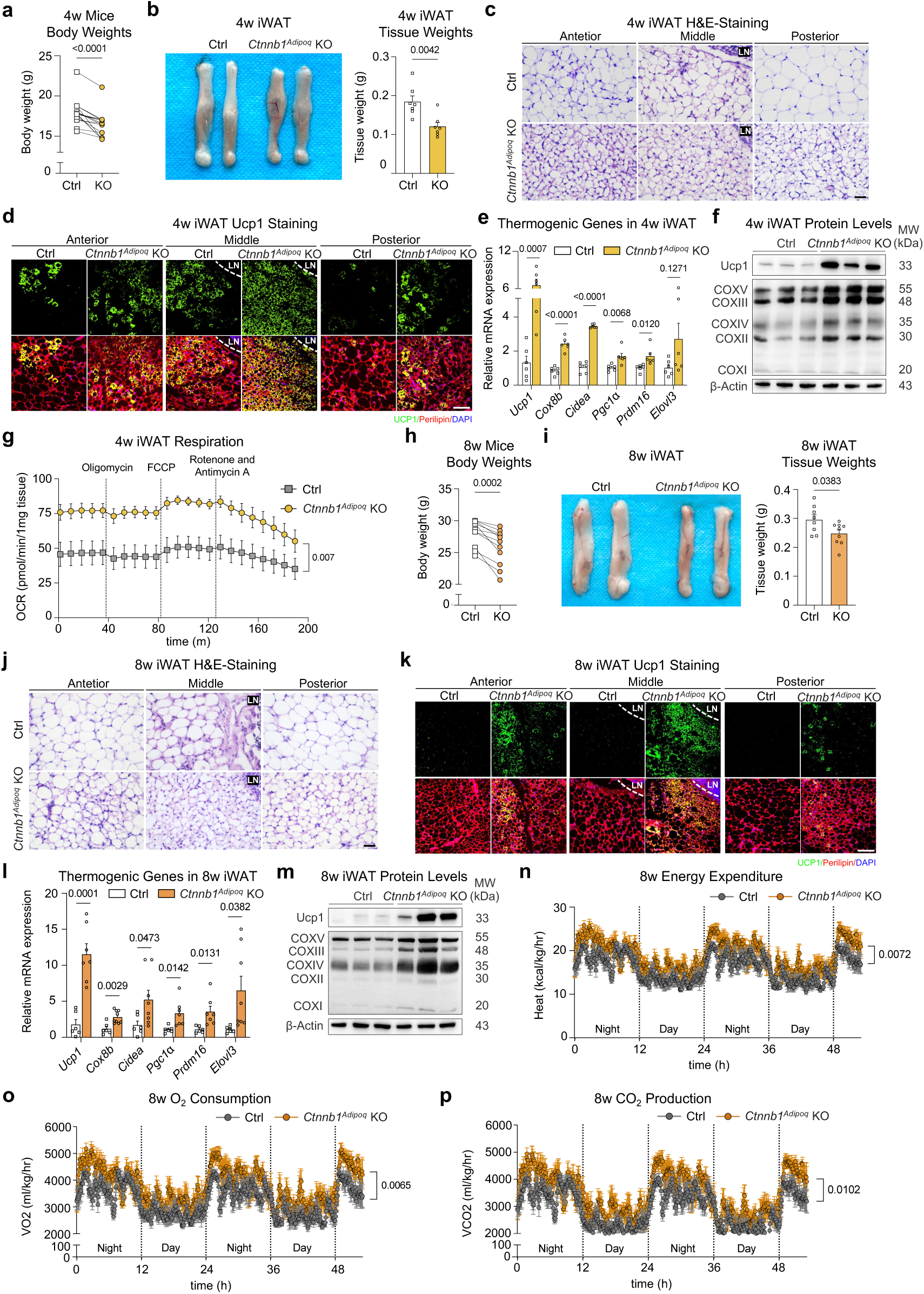
Adipocyte β-catenin ablation enhances developmental beiging in iWAT. **a** Body weight of 4-week-old *Adipoq*^Cre^;*Ctnnb1*^flox/flox^ (*Ctnnb1^Adipoq^* KO) and control (Ctrl) littermates (*n* = 8 *Adipoq*^Cre^;*Ctnnb1*^flox/flox^, *n* = 9 control). **b** iWAT macroscopic appearance and iWAT mass of 4-week-old *Adipoq*^Cre^;*Ctnnb1*^flox/flox^ and control littermates (*n* = 7). **c** Representative H&E-stained iWAT sections in (**b**) (*n* = 3). White text LN indicates lymph node. Scale bar, 50 μm. **d** Immunofluorescent staining in (**b**) (*n* = 3). White dashed line outlines LN. DAPI, 4,6-diamidino-2-phenylindole. Scale bar, 100 μm. **e** mRNA expression of thermogenic genes in (**b**) (*n* = 6). **f** Immunoblotting for Ucp1 and OXPHOS in (**b**) (*n* = 3). **g** Oxygen consumption rate (OCR) plots in (**b**) (*n* = 3). **h** Body weight of 8-week-old *Adipoq*^Cre^;*Ctnnb1*^flox/flox^ and control littermates (*n* = 10 *Adipoq*^Cre^;*Ctnnb1*^flox/flox^, *n* = 8 control). **i** iWAT macroscopic appearance and iWAT mass of 8-week-old *Adipoq*^Cre^;*Ctnnb1*^flox/flox^ and control littermates (*n* = 8 *Adipoq*^Cre^;*Ctnnb1*^flox/flox^, *n* = 9 control). **j** Representative H&E-stained iWAT sections in (**i**) (*n* = 3). White text LN indicates lymph node. Scale bar, 50 μm. **k** Immunofluorescent staining in (**i**) (*n* = 3). White dashed line outlines LN. Scale bar, 100 μm. **l** mRNA expression of thermogenic genes in (**i**) (*n* = 7-9 *Adipoq*^Cre^;*Ctnnb1*^flox/flox^, *n* = 6 control). **m** Immunoblotting for Ucp1 and OXPHOS in (**i**) (*n* = 3). **n–p** Energy expenditure (**n**), whole-body oxygen (O_2_) consumption (**o**), carbon dioxide (CO_2_) production (**p**) measured in metabolic cages of 8-week-old *Adipoq*^Cre^;*Ctnnb1*^flox/flox^ and control mice (*n* = 5). The levels of mRNA expression are normalized to that of *36B4*. Data are the mean ± s.e.m. Statistical differences were performed using two-way ANOVA followed by Tukey’s test or paired two-sided Student’s *t*-test or unpaired two-sided Student’s *t*-test.

Because β-catenin has well-established structural roles in cell-cell adhesion and cytoskeletal organization in addition to its function as a transcriptional effector of the canonical Wnt signaling^31,32^, we next asked whether the thermogenic phenotype observed in *Adipoq*^Cre^;*Ctnnb1*^flox/flox^ mice reflects loss of Wnt/β-catenin transcriptional outputs or disruption of these structural functions. To functionally separate these roles, we used a transcription-specific *Ctnnb1* mutant allele (*Ctnnb1*^dm^), in which mutations in the N- and C-terminal domains abolish its binding to transcriptional co-activators while preserving β-catenin’s junctional structural functions^33^. We generated *Adipoq*^Cre^;*Ctnnb1*^dm/flox^ mice by crossing *Adipoq*^Cre^;*Ctnnb1*^flox/+^ allele with *Ctnnb1*^dm/+^ mice (Supplementary Fig. 3e), such that adipocytes express a truncated β-catenin protein that is structurally intact but transcriptionally inactive in the canonical Wnt pathway. We found that thermogenic genes in iWAT from *Adipoq*^Cre^;*Ctnnb1*^dm/flox^ mice were consistently upregulated (Supplementary Fig. 3f), similar to the changes observed in *Adipoq*^Cre^;*Ctnnb1*^flox/flox^ mice, supporting the view that the thermogenic phenotype caused by adipocyte β-catenin depletion is primarily attributable to the loss of Wnt/β-catenin signaling.

Given that UCP1 expression in iWAT becomes detectable from approximately postnatal day 14^24^, we used *Ucp1*^Cre^ to further exclude the possibility that these effects were driven by early alterations in adipose development. We therefore generated *Ucp1*^Cre^;*Ctnnb1*^flox/flox^ mice in which *Ctnnb1* was conditionally ablated in *Ucp1*-expressing beige adipocytes (Supplementary Fig. 3h). In such mice, blockade of Wnt/β-catenin produced changes similar to those observed in *Adipoq*^Cre^;*Ctnnb1*^flox/flox^ mice, including reduced body weight and iWAT mass, decreased adipocyte size, and increased thermogenic gene and protein expression (Supplementary Fig. 3i–m), consistent with a previous report^34^. Together with the *Adipoq*^Cre^;*Ctnnb1*^flox/flox^, *Adipoq*^Cre^;*Ctnnb1*^dm/flox^, and *Ucp1*^Cre^;*Ctnnb1*^flox/flox^ models, these results indicate a primary role for Wnt/β-catenin signaling in restraining thermogenic function in mature adipocytes.

### Adipocyte β-catenin ablation selectively enhances thermogenesis in iWAT

To determine whether enhanced thermogenesis in *Adipoq*^Cre^;*Ctnnb1*^flox/flox^ iWAT could be explained by altered adipocyte differentiation, we examined adipogenesis in vivo and in vitro. In iWAT, adipogenic markers, assessed by RT-qPCR and immunoblotting, were not substantially reduced in *Adipoq*^Cre^;*Ctnnb1*^flox/flox^ mice compared with controls (Supplementary Fig. 4a,b), and *Tle3*, a transcriptional co-regulator implicated in maintaining white adipocyte gene program^35,36^, was expressed at similar levels in both genotypes (Supplementary Fig. 4c). Likewise, adipocytes differentiated from *Adipoq*^Cre^;*Ctnnb1*^flox/flox^ and control SVF cells exhibited comparable expression of adipogenic markers (Supplementary Fig. 4d,e). To test if lipogenic capacity was compromised in *Adipoq*^Cre^;*Ctnnb1*^flox/flox^ iWAT, given the smaller fat cell size, we examined the expression of key genes involved in de novo lipid synthesis, including *Srebp1*, *Mlxipl*, *Acc1*, *Scd1*, *Fasn*, *Acly*, and *Dgat*. Their expression levels were identical between *Adipoq*^Cre^;*Ctnnb1*^flox/flox^ and control mice, indicating no major defect in lipogenesis (Supplementary Fig. 4f). In addition, to evaluate endocrine function and systemic metabolic status in developmental *Adipoq*^Cre^;*Ctnnb1*^flox/flox^ mice, serum ELISA analyses were performed. These assays revealed no significant differences in adipokines such as adiponectin, or in standard circulating metabolic parameters, between *Adipoq*^Cre^;*Ctnnb1*^flox/flox^ and control mice (Supplementary Fig. 4g). These data indicate that adipocyte-specific β-catenin deletion primarily promotes thermogenic capacity in adipose tissue.

Because *Adipoq*^Cre^;*Ctnnb1*^flox/flox^ mice elicited an enhanced beiging response in subcutaneous WAT, we next asked whether other adipose depots were similarly affected. We therefore examined interscapular BAT, the classical thermogenic adipose organ in which brown adipocytes mediate non-shivering thermogenesis^2^, in control and β-catenin-deficient cohorts. BAT mass, brown adipocyte morphology, and thermogenic/adipogenic marker expression were broadly comparable between control and *Adipoq*^Cre^;*Ctnnb1*^flox/flox^ mice at both 4 and 8 weeks, and were likewise unchanged in 4-week *Ucp1*^Cre^;*Ctnnb1*^flox/flox^ mice relative to controls (Supplementary Fig. 5a–j). We also assessed eWAT, a visceral white fat depot where beige recruitment is typically limited^5^. In eWAT from 4- and 8-week *Adipoq*^Cre^;*Ctnnb1*^flox/flox^ and control mice, expression of thermogenic and adipogenic genes did not differ significantly between genotypes (Supplementary Fig. 5k,l). Thus, these findings suggest that adipocyte Wnt/β-catenin signaling acts in a depot-selective manner to constrain beige adipocyte recruitment and adaptive thermogenesis in iWAT.

### Wnt/β-catenin suppression sustains beige fat thermogenesis under ambient conditions into adulthood and increased energy expenditure in adult mice

To examine whether the enhanced thermogenesis persists into adulthood, we analyzed 8-week-old *Adipoq*^Cre^;*Ctnnb1*^flox/flox^ and control mice maintained at room temperature. We found that adult *Adipoq*^Cre^;*Ctnnb1*^flox/flox^ mice continued to exhibit reduced body weight, fat mass, and adipocyte size in iWAT compared with controls (Fig. 2h,i, Supplementary Fig. 6a). These morphological changes were accompanied by sustained upregulation of thermogenic genes and protein in iWAT, whereas adipogenic markers showed no appreciable reduction in *Adipoq*^Cre^;*Ctnnb1*^flox/flox^ mice (Fig. 2j–m, Supplementary Fig. 6b–f). Consistent with a transcriptional mechanism, thermogenesis-related gene expression in iWAT from adult *Adipoq*^Cre^;*Ctnnb1*^dm/flox^ mice was also significantly increased (Supplementary Fig. 6g). To further explore if the durable beiging translates into enhanced systemic metabolism, we measured the rates of whole-body energy expenditure (EE). *Adipoq*^Cre^;*Ctnnb1*^flox/flox^ mice had a higher level of EE, oxygen consumption, and carbon dioxide production relative to their controls (Fig. 2n–p), with no differences in food intake or respiratory exchange ratio (RER) (Supplementary Fig. 6h,i). These findings provide evidence that inhibition of Wnt/β-catenin signaling in adipocytes leads to a persistent elevated iWAT adaptive thermogenesis and whole-body energy expenditure into adulthood.

### Reduced Wnt/β-catenin signaling engages a non-canonical Wnt/Ca^2+^-AMPK axis in developmental iWAT beiging

To elucidate the molecular mechanisms by which β-catenin deletion enhances beige adipocyte thermogenesis during peri-weaning period, we ran bulk RNA-seq on iWAT from 4-week-old *Adipoq*^Cre^;*Ctnnb1*^flox/flox^ and control mice. As expected, volcano plots revealed upregulation of thermogenesis-related genes in *Adipoq*^Cre^;*Ctnnb1*^flox/flox^ iWAT (Supplementary Fig. 7a). Notably, functional enrichment analysis showed that the transcriptional changes were significantly enriched in pathways related to PPAR signaling, Ca^2+^ signaling, and AMPK signaling (Fig. 3a), all of which have established roles in promoting oxidative metabolism in thermogenic adipocytes^12,15,16,37,38^. On the basis of these observations and given the well-known antagonistic relationship between the canonical Wnt and non-canonical Wnt/Ca^2+^ signaling^39–44^, we reasoned that β-catenin loss might shift signaling towards the non-canonical Wnt/Ca^2+^ pathway that fits into Ca^2^⁺-sensitive AMPK-PPAR oxidative programs. To test this hypothesis, we subsequently examined the non-canonical Wnt ligands and their receptors in wild-type iWAT across developmental and adult stages. Among these candidates, the ligand *Wnt5a* and its receptor *Fzd5* showed significantly higher expression in 4-week iWAT than in 8-week iWAT (Fig. 3b). Concomitantly, immunoblotting revealed increased phosphorylation of CaMKK2 and AMPK, together with elevated Wnt5a, Sirt1 and Pgc1α protein abundance, in peri-weaning compared with adult iWAT (Fig. 3c, Supplementary Fig. 7b,c), consistent with activation of a Wnt5a-Ca^2+^-AMPK cascade and induction of downstream Sirt1-Pgc1α oxidative programs during developmental beige adipocyte recruitment^10,45^. Given that AMPK is an energy-sensing kinase that promotes triglyceride breakdown in adipocytes^12^, we then examined the key lipolytic enzyme Atgl together with Pparγ, a PPAR family transcription factor that links lipid-derived signals to oxidative gene expression^15,16^. We found that the expression of both Atgl and Pparγ was higher in peri-weaning than in adult iWAT (Fig. 3c, Supplementary Fig. 7d), suggesting engagement of an AMPK-linked lipolytic and PPAR-associated oxidative component.

**Fig. 3.**
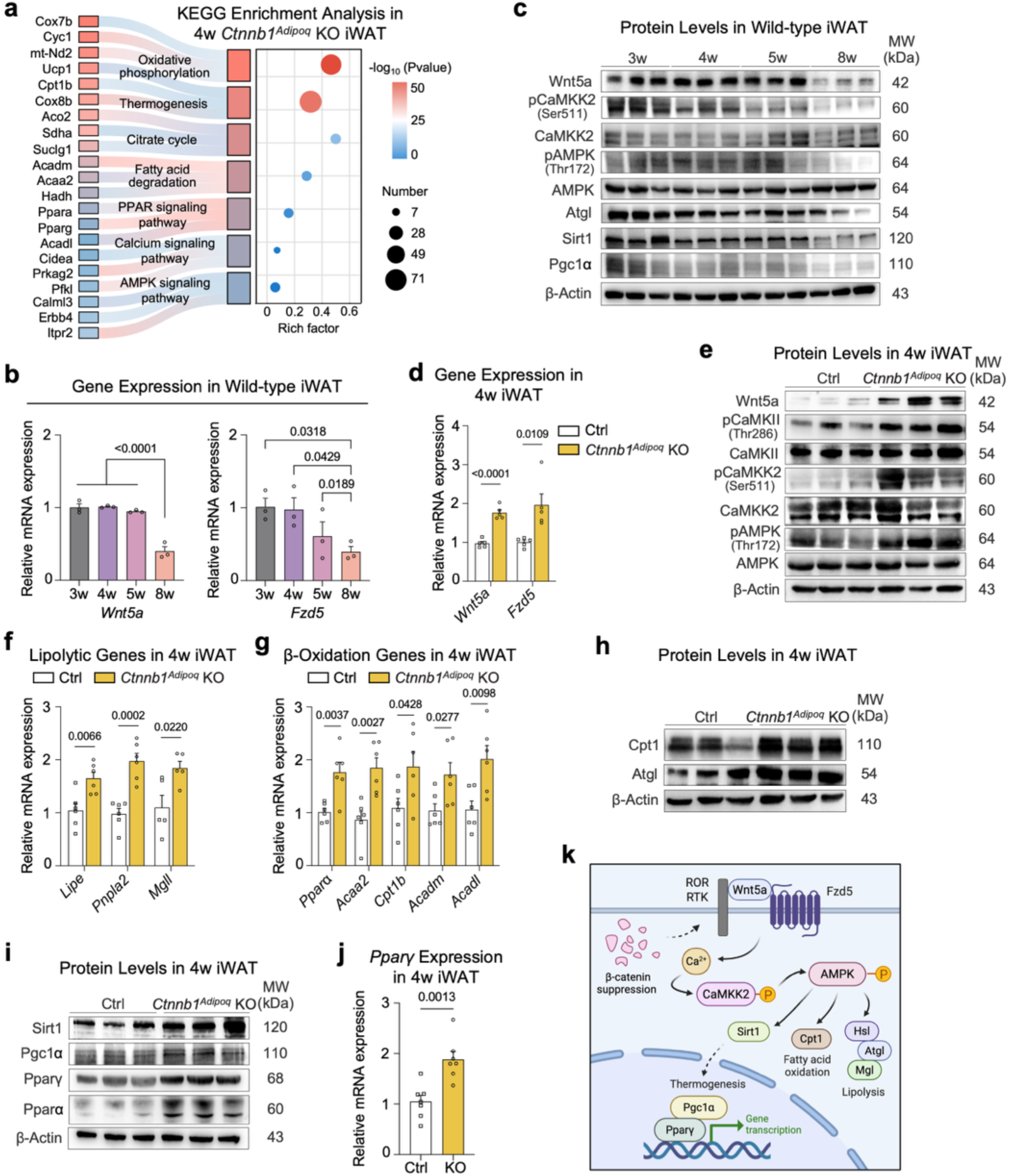
Reduced Wnt/β-catenin signaling engages a non-canonical Wnt/Ca^2+^-AMPK axis in developmental iWAT beiging. **a** RNA-seq was performed in iWAT of 4-week-old *Adipoq*^Cre^;*Ctnnb1*^flox/flox^ and control mice (*n* = 3), with a threshold of |FC| > 1.5 and adjusted P-value < 0.05. KEGG enrichment analysis of genes upregulated in *Adipoq*^Cre^;*Ctnnb1*^flox/flox^ mice relative to control mice. **b** mRNA expression of *Wnt5a* and *Fzd5* in iWAT of wild-type male mice at 3, 4, 5, and 8 weeks of age (*n* = 3). **c** Immunoblotting for Wnt5a, pCaMKK2, CaMKK2, pAMPK, AMPK, Atgl, Sirt1, Pgc1α in iWAT of wild-type male mice at 3, 4, 5, and 8 weeks of age (*n* = 3). **d** *Wnt5a* and *Fzd5* expression in iWAT of 4-week-old *Adipoq*^Cre^;*Ctnnb1*^flox/flox^ and control mice (*n* = 3). **e** Immunoblotting for Wnt5a, pCaMKII, CaMKII, pCaMKK2, CaMKK2, pAMPK, AMPK in iWAT of 4-week-old *Adipoq*^Cre^;*Ctnnb1*^flox/flox^ and control mice (*n* = 3). **f** mRNA expression of triglyceride lipolysis genes in iWAT of 4-week-old *Adipoq*^Cre^;*Ctnnb1*^flox/flox^ and control mice (*n* = 5–6). **g** mRNA expression of fatty acid β-oxidation genes in iWAT of 4-week-old *Adipoq*^Cre^;*Ctnnb1*^flox/flox^ and control mice (*n* = 6). **h** Immunoblotting for Cpt1, Atgl in iWAT of 4-week-old *Adipoq*^Cre^;*Ctnnb1*^flox/flox^ and control mice (*n* = 3). **i** Immunoblotting for Sirt1, Pgc1α, Pparγ, Pparɑ in iWAT of 4-week-old *Adipoq*^Cre^;*Ctnnb1*^flox/flox^ and control mice (*n* = 3). **j** mRNA expression of *Pparγ* in iWAT of 4-week-old *Adipoq*^Cre^;*Ctnnb1*^flox/flox^ and control mice (*n* = 7). **k** Schematic summary of β-catenin suppression results in activation of Wnt/Ca^2+^-AMPK axis, highlighting non-canonical pathways involved in the thermogenic response. The levels of mRNA expression are normalized to that of *36B4*. Data are the mean ± s.e.m. Statistical analyses used were one-way ANOVA with Tukey’s correction for multiple comparisons or unpaired two-tailed *t*-tests.

We next asked if this Wnt5a-Ca^2^⁺-AMPK axis is further amplified upon adipocyte β-catenin deletion. In 4-week-old *Adipoq*^Cre^;*Ctnnb1*^flox/flox^ mice, *Wnt5a* and *Fzd5* expression in iWAT was increased (Fig. 3d), while remaining unchanged in BAT and eWAT (Supplementary Fig. 7e), underlying the depot-specific phenotype. Consistently, phosphorylation of CaMKII, CaMKK2, and AMPK was markedly elevated compared with controls (Fig. 3e, Supplementary Fig. 7f), indicating robust activation of this axis in the absence of β-catenin. We therefore examined lipid catabolic and oxidative programs in *Adipoq*^Cre^;*Ctnnb1*^flox/flox^ iWAT. *Adipoq*^Cre^;*Ctnnb1*^flox/flox^ mice had a significantly higher expression of mRNAs and proteins involved in triglyceride lipolysis and fatty acid β-oxidation (Fig. 3f–h, Supplementary Fig. 7g). In addition, adipocyte *Ctnnb1* knockout at the developmental stage led to significantly increased expression of Sirt1 and Pgc1α, as well as Pparγ and Pparα in iWAT (Fig. 3i,j, Supplementary Fig. 7h), indicative of an inhibitory role of Wnt/β-catenin signaling in constraining AMPK-mediated developmental beiging.

Because the metabolic benefits of developmental β-catenin loss are preserved into adulthood, we also interrogated this axis in 8-week-old *Adipoq*^Cre^;*Ctnnb1*^flox/flox^ mice. Interestingly, we observed that *Wnt5a* and *Fzd5* mRNA levels were no longer significantly different between genotypes (Supplementary Fig. 8a), although phosphorylation of CaMKK2 and AMPK and expression of genes related to fatty acid oxidation and lipolysis remained elevated in *Adipoq*^Cre^;*Ctnnb1*^flox/flox^ iWAT (Supplementary Fig. 8j–o). This pattern suggests that transcriptional induction of the non-canonical Wnt/Ca^2^⁺ signaling is most prominent during the peri-weaning window, whereas a higher Ca^2^⁺-AMPK activity state and its associated oxidative-lipolytic activities sustained in *Adipoq*^Cre^;*Ctnnb1*^flox/flox^ iWAT at a later adult stage. Together, these findings support a model in which reduced Wnt/β-catenin signaling in adipocytes activates the non-canonical Wnt/Ca^2+^-AMPK pathway, with downstream Sirt1-Pgc1α-PPAR-dependent oxidative and lipolytic outputs that contribute to peri-weaning beige adipocyte thermogenesis (Fig. 3k).

### β-catenin inhibition elicits Ca^2^⁺-dependent thermogenic activation in cultured peri-weaning and adult adipocytes

To extend these findings to an in vitro setting, SVF cells were isolated from iWAT of 4-week-old wild-type mice, differentiated into mature adipocytes, and then treated with the β-catenin destabilizer MSAB for 4 days^46^. In parallel, rosiglitazone was included as a positive beiging control^47^ from the onset of differentiation until sample collection (Fig. 4a). Compared with DMSO-treated cells, both MSAB and rosiglitazone administration yielded adipocytes with reduced lipid droplet size and increased droplet number, resulting in a more multilocular morphology characteristic of beige adipocytes (Fig. 4b). Immunoblotting confirmed that MSAB lowered β-catenin protein abundance and attenuated Wnt/β-catenin signaling activity (Supplementary Fig. 9a,b), whereas adipogenic marker expression was largely unchanged across conditions (Supplementary Fig. 9c,d). In agreement with the in vivo phenotype, MSAB alone increased thermogenic marker expression, and the combination of MSAB and rosiglitazone produced the strongest induction (Fig. 4c). Analysis of signaling components further showed that the suppression of Wnt/β-catenin was accompanied by higher Wnt5a levels, increased phosphorylation of CaMKK2 and AMPK, and elevated Sirt1, Pgc1α, Pparγ, and Ucp1 proteins (Fig. 4d, Supplementary Fig. 9e,f), recapitulating the Ca^2+^-AMPK-PPAR-related thermogenic signature observed in vivo.

**Fig. 4.**
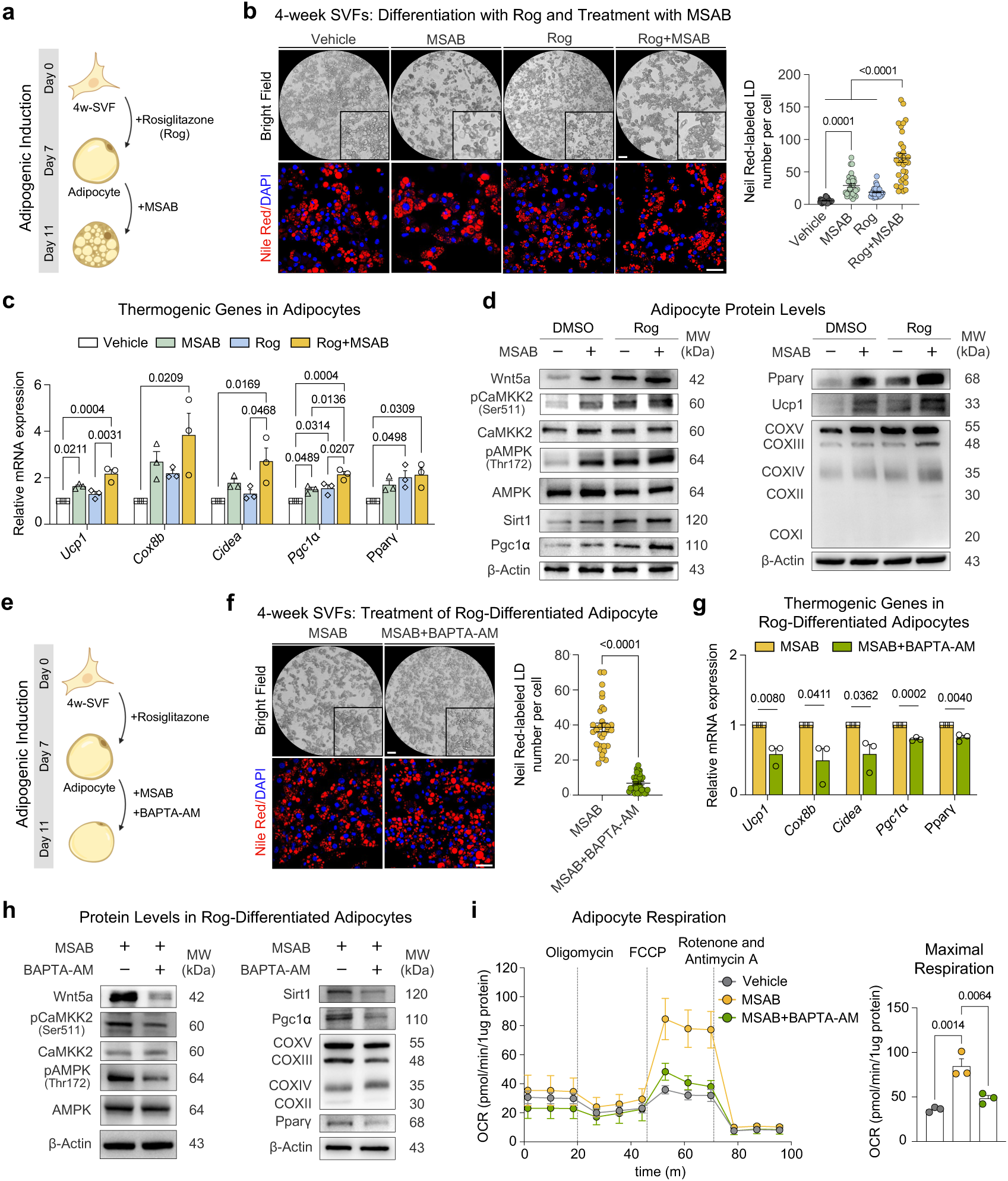
β-catenin suppression elicits Ca2⁺-dependent thermogenic activation in cultured peri-weaning adipocytes. **a** Experimental strategy to suppress β-catenin in vitro by employing adipocytes differentiated from SVFs of 4-week-old wild-type mouse iWAT. **b** Microscopy images of adipocytes following treatment with vehicle (0.1% DMSO), MSAB, Rosiglitazone, or MSAB plus Rosiglitazone (*n* = 3 biological replicates). Scale bar, 200 μm. Nile Red and DAPI were used to stain lipid droplets and nuclei, respectively (*n* = 3 biological replicates). Scale bar, 20 μm. The right panel displays the statistical analysis of Nile Red-labeled lipid droplets number per cell (*n* = 32 cells). **c** mRNA expression of thermogenic genes in (**b**) (*n* = 3 biological replicates). **d** Immunoblotting for Wnt5a/Ca²⁺-AMPK-PPAR axis and thermogenic protein in (**b**) (*n* = 3 biological replicates). **e** Experimental strategy to suppress β-catenin while inhibiting the Ca²⁺ pathway in vitro by employing adipocytes differentiated from the SVFs of 4-week-old wild-type mouse iWAT. **f** Microscopy images of Rosiglitazone-differentiated adipocytes treated with vehicle (0.1% DMSO), MSAB, or MSAB plus BAPTA-AM (*n* = 3 biological replicates). Scale bar, 200 μm. Nile Red and DAPI were used to stain lipid droplets and nuclei, respectively (*n* = 3 biological replicates). Scale bar, 20 μm. The right panel displays the statistical analysis of Nile Red-labeled lipid droplets number per cell (*n* = 32 cells). **g** mRNA expression of thermogenic genes in (**f**) (*n* = 3 biological replicates). **h** Immunoblotting for Wnt5a/Ca²⁺-AMPK-PPAR axis and thermogenic protein in (**f**) (*n* = 3 biological replicates). **i** OCR plots and measured maximal respiration levels across the three treatment groups (Vehicle, MSAB, MSAB and BAPTA-AM) (*n* = 3 cells). The levels of mRNA expression are normalized to that of *36B4*. Data are the mean ± s.e.m. Statistical analyses used were unpaired two-sided Student’s *t*-tests or two-way ANOVA followed by Tukey’s multiple-comparisons test or one-way ANOVA with Tukey’s correction for multiple comparisons.

To test whether Ca^2+^ is required for the thermogenic effects of Wnt/β-catenin inhibition, we added the intracellular Ca^2+^ chelator BAPTA-AM^48,49^ on top of MSAB and rosiglitazone treatment (Fig. 4e). Supplementation with the Ca^2^⁺ inhibitor shifted adipocyte morphology toward larger and more unilocular lipid droplets despite comparable adipogenic marker expression (Fig. 4f, Supplementary Fig. 9g,h). Meanwhile, thermogenic marker expression was significantly compromised, and Wnt5a-, CaMKK2- and AMPK-related readouts were attenuated (Fig. 4g,h, Supplementary Fig. 9i). BAPTA-AM administration also tended to increase expression of Wnt/β-catenin target genes such as *Axin2*, *Myc*, and *Lef1* (Supplementary Fig. 9j), in line with reciprocal interactions between Ca^2^⁺-dependent and canonical Wnt pathways. Functionally, Seahorse assay showed that maximum respiratory capacity was highest in MSAB-treated adipocytes and was remarkably blunted by Ca^2^⁺ chelation (Fig. 4i). These data indicate that Ca^2^⁺-dependent signaling is necessary for the full thermogenic response to Wnt/β-catenin inhibition in adipocytes.

Given the age-dependent decline in spontaneous beiging potential and the concomitant increase in Wnt/β-catenin activity in vivo, we next asked if pharmacological suppression of Wnt/β-catenin signaling pathway in vitro could restore a thermogenic phenotype in adult adipocytes. To this end, SVF cells from iWAT of 8-week-old wild-type mice were differentiated and subjected to the same MSAB and BAPTA-AM treatments. In this adult setting, Wnt/β-catenin inhibition again induced a beige-like morphology and upregulated thermogenic markers (Supplementary Fig. 10a–c), whereas concomitant Ca^2^⁺ chelation attenuated these effects (Supplementary Fig. 10d–f). Thus, adipocytes derived from both developmental and adult iWAT retain an intrinsic capacity to increase thermogenesis in response to Wnt/β-catenin inhibition, which depends on intact Ca^2^⁺ signaling. These findings further support Wnt/β-catenin activity as a determinant of age-dependent divergence in adipocyte thermogenic capacity.

### Non-canonical Wnt5a-Ca^2+^ signaling supports adipocyte beiging only when canonical Wnt/β-catenin is suppressed

To dissect the contribution of non-canonical Wnt/Ca^2+^ signaling to the beiging response induced by β-catenin inhibition in vitro, iWAT-derived SVF cells from 4-week-old wild-type mice were differentiated into adipocytes and then treated for 4 days with Box5, a selective Wnt5a/Ca^2+^ antagonist^50^, in the presence or absence of Wnt/β-catenin signaling inhibitor MSAB. (Fig. 5a). In MSAB-treated adipocytes, the addition of Box5 resulted in relatively larger lipid droplets and loss of the bead-like, multilocular morphology observed with MSAB alone (Fig. 5b). Consistent with these morphological changes, co-treatment with MSAB and Box5 markedly blunted induction of the thermogenic program, as reflected by reduced expression of Ucp1 and other beige markers at both mRNA and protein levels, and attenuated Wnt5a-Ca^2+^-AMPK signaling activity, without substantially altering adipogenic marker expression (Fig. 5c,d, Supplementary Fig. 11a–c). By contrast, in the absence of MSAB, Box5 alone had minimal effects on lipid droplet morphology, thermogenic markers or Wnt5a-Ca^2^⁺-AMPK signaling compared with vehicle-treated cells, suggesting that perturbation of Wnt5a-Ca^2+^ signaling does not appreciably influence adipocyte beiging when Wnt/β-catenin activity is intact (Fig. 5b–d, Supplementary Fig. 11a).

**Fig. 5.**
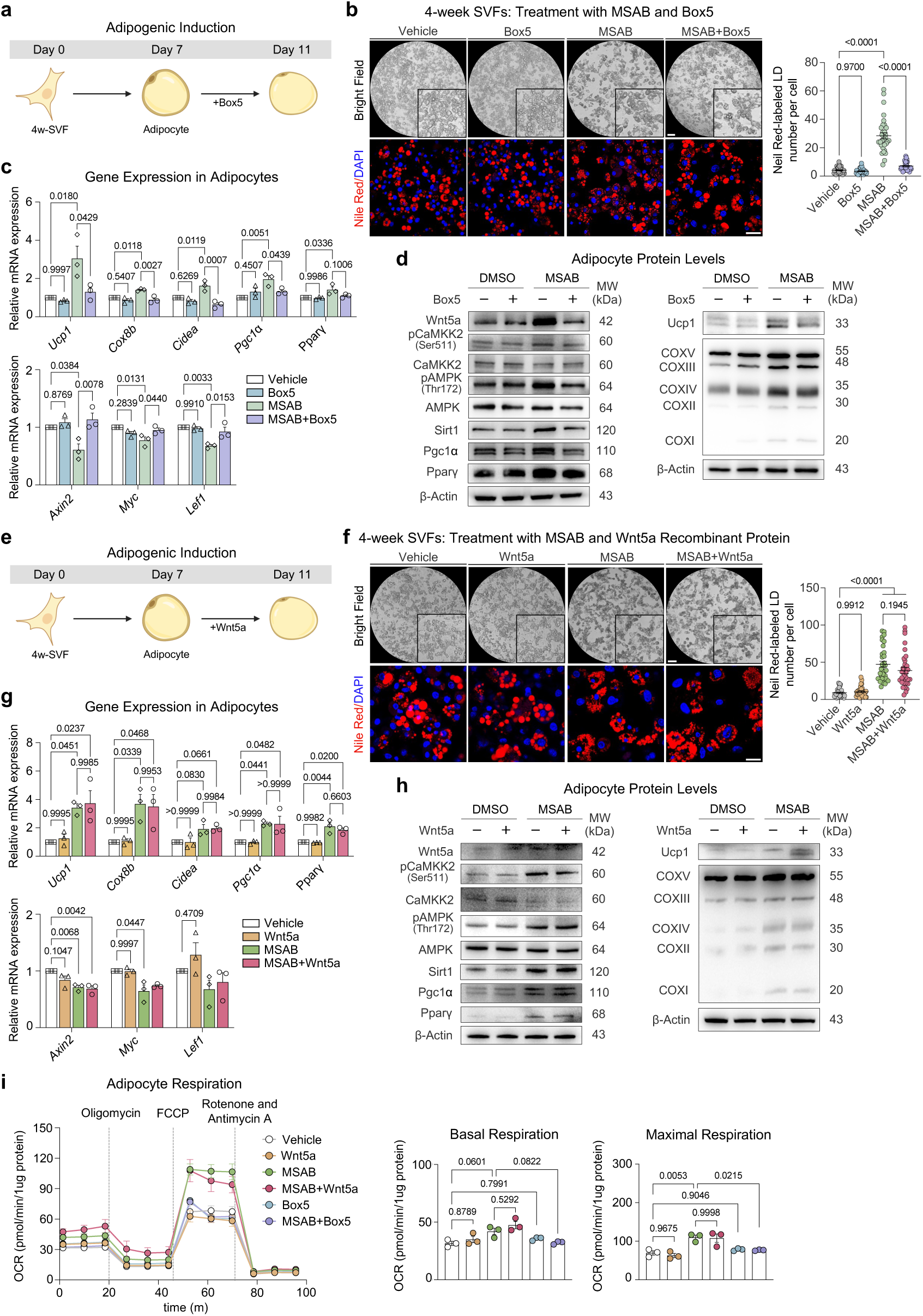
Imbalance between canonical and non-canonical Wnt pathways drives beige formation. **a** Experimental strategy to suppress Wnt5a/Ca²⁺ pathway in vitro by employing adipocytes differentiated from the SVFs of 4-week-old wild-type mouse iWAT. **b** Microscopy images of adipocytes following treatment with vehicle (0.1% DMSO), Box5, MSAB, or MSAB plus Box5 (*n* = 3 biological replicates). Scale bar, 200 μm. Nile Red and DAPI were used to stain lipid droplets and nuclei, respectively (*n* = 3 biological replicates). Scale bar, 20 μm. The right panel displays the statistical analysis of Nile Red-labeled lipid droplets number per cell (*n* = 32 cells). **c** mRNA expression of thermogenic genes and downstream genes of Wnt/β-catenin signaling pathway in (**b**) (*n* = 3 biological replicates). **d** Immunoblotting for Wnt5a/Ca²⁺-AMPK-PPAR axis and thermogenic protein in (**b**) (*n* = 3 biological replicates). **e** Experimental strategy to supply exogenous Wnt5a recombinant protein in vitro by employing adipocytes differentiated from the SVFs of 4-week-old wild-type mouse iWAT. **f** Microscopy images of adipocytes following treatment with vehicle (0.1% DMSO), Wnt5a recombinant protein, MSAB, or MSAB plus Wnt5a recombinant protein (*n* = 3 biological replicates). Scale bar, 200 μm. Nile Red and DAPI were used to stain lipid droplets and nuclei, respectively (*n* = 3 biological replicates). Scale bar, 20 μm. The right panel displays the statistical analysis of Nile Red-labeled lipid droplets number per cell (*n* = 32 cells). **g** mRNA expression of thermogenic genes and downstream genes of Wnt/β-catenin signaling pathway in (**f**) (*n* = 3 biological replicates). **h** Immunoblotting for Wnt5a/Ca²⁺-AMPK-PPAR axis and thermogenic protein in (**f**) (*n* = 3 biological replicates). **i** OCR plots, measured basal respiration and measured maximal respiration levels in cultured peri-weaning adipocytes across the six treatment groups (Vehicle, Wnt5a recombinant protein, MSAB, MSAB plus Wnt5a recombinant protein, Box5, MSAB plus Box5) (*n* = 3 cells). The levels of mRNA expression are normalized to that of *36B4*. Data are the mean ± s.e.m. Statistical analyses used were two-way ANOVA followed by Tukey’s multiple-comparisons test or one-way ANOVA with Tukey’s correction for multiple comparisons.

We next asked whether activation of Wnt5a signaling is sufficient to promote adipocyte beiging. Differentiated adipocytes were treated for 4 days with recombinant Wnt5a, with or without concomitant MSAB treatment (Fig. 5e). We observed that adipogenic capacity was similar across groups, but a bead-like, multilocular morphology emerged only when Wnt/β-catenin was inhibited; Wnt5a alone did not appreciably alter lipid droplet morphology or adipogenesis (Fig. 5f, Supplementary Fig. 11d). Consistently, recombinant Wnt5a in the absence of MSAB failed to activate Wnt5a-Ca^2+^-AMPK signaling or to induce a significant beiging response. (Fig. 5g,h, Supplementary Fig. 11e). Measurements of oxygen consumption showed that respiratory capacity was increased in MSAB and MSAB plus Wnt5a groups, but this enhancement was diminished when Wnt5a pathway was blocked by Box5, whereas Wnt5a or Box5 alone were largely ineffective (Fig. 5i). Notably, exogenous Wnt5a did not further augment beiging beyond that achieved by MSAB alone, suggesting that relief of Wnt/β-catenin signaling rather than Wnt5a availability is rate-limiting under these conditions. Collectively, these results indicate that Wnt5a-responsive Ca^2+^ signaling reinforces adipocyte thermogenesis when Wnt/β-catenin activity is lowered, but is not sufficient to trigger beiging when the canonical Wnt signaling remains active.

### Wnt/β-catenin inhibition promotes thermogenic activation in human subcutaneous adipocytes

To determine whether this mechanism is conserved in humans, we performed in vitro studies using human subcutaneous adipocytes obtained from liposuction aspirates. SVF cells were differentiated to maturity and then treated for 4 days with MSAB to inhibit Wnt/β-catenin signaling (Fig. 6a). Similar to murine adipocytes, MSAB-treated human adipocytes exhibited a highly reduction in lipid droplet size and a more punctate, multilocular appearance without altering adipogenic marker expression (Fig. 6b,c). Immunoblotting and gene expression analyses showed that inhibition of Wnt/β-catenin also gave rise to increased WNT5A expression, enhanced CAMKK2 and AMPK phosphorylation, and upregulated SIRT1, PGC1α and thermogenic markers (Fig. 6d,e). These findings indicate that suppression of Wnt/β-catenin signaling can trigger a Wnt5a-Ca^2+^-AMPK-linked thermogenic response in human subcutaneous adipocytes, consistent with the mechanism defined in mice.

**Fig. 6.**
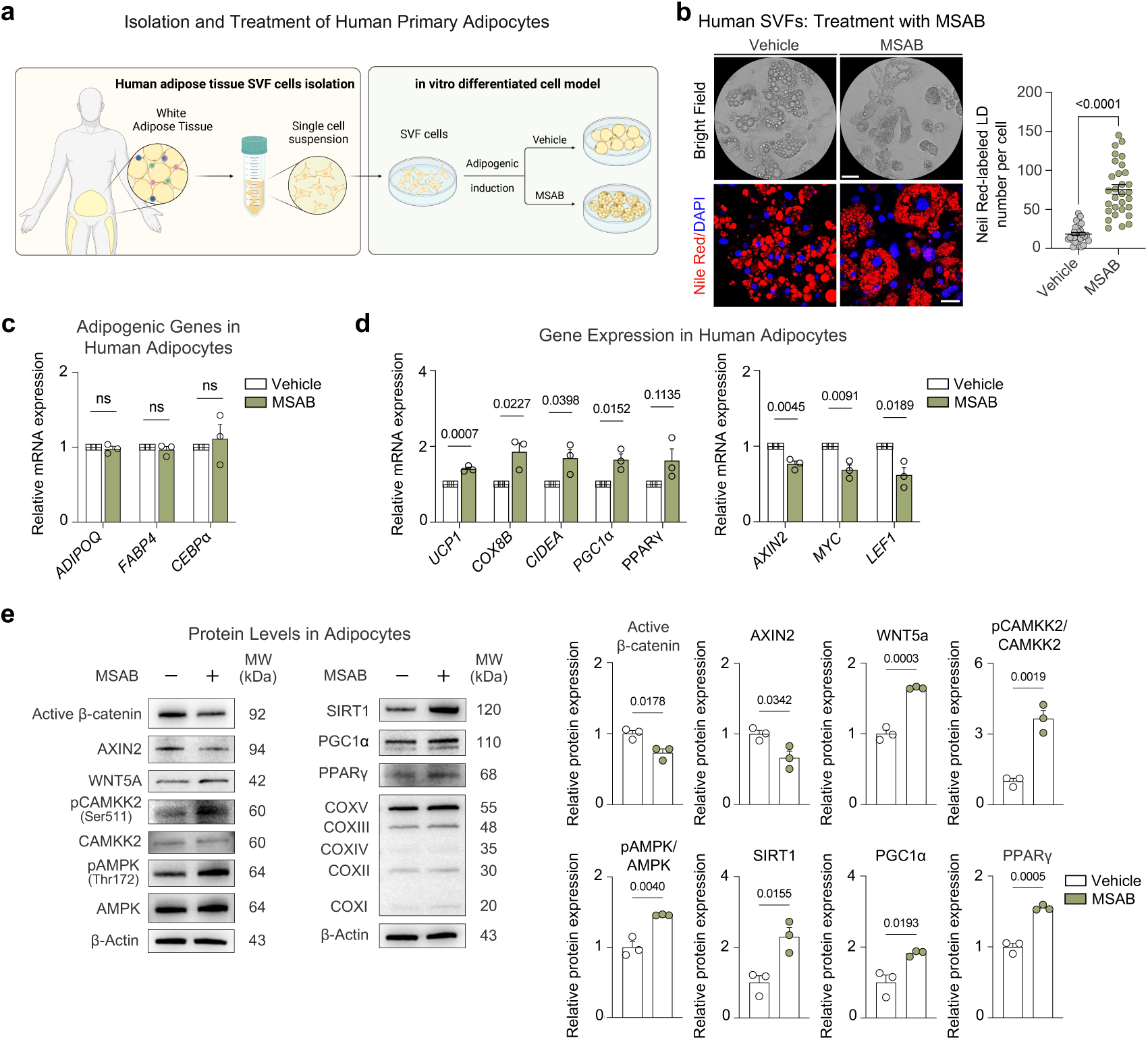
β-catenin reduction promotes thermogenic activation in human subcutaneous adipocytes. **a** Experimental strategy to suppress β-catenin using adipocytes differentiated from human subcutaneous lipoaspirates. **b** Microscopy images of human subcutaneous adipocytes following treatment with vehicle (0.1% DMSO) and MSAB after maturation (*n* = 3 biological replicates). Scale bar, 200 μm. Nile Red and DAPI were used to stain lipid droplets and nuclei, respectively (*n* = 3 biological replicates). Scale bar, 30 μm. The right panel displays the statistical analysis of Nile Red-labeled lipid droplets number per cell (*n* = 30 cells). **c** mRNA expression of adipogenic genes in (**b**) (*n* = 3 biological replicates). **d** mRNA expression of thermogenic genes and downstream genes of Wnt/β-catenin signaling pathway in (**b**) (*n* = 3 biological replicates). **e** Immunoblotting for Wnt5a/Ca²⁺-AMPK-PPAR axis, Wnt/β-catenin signaling pathway and thermogenic protein in (**b**) (*n* = 3 biological replicates). The levels of mRNA expression are normalized to that of *36B4*. Data are the mean ± s.e.m. Statistical analyses used were unpaired two-sided Student’s *t*-tests.

## Discussion

Beige adipocytes have emerged as attractive targets for improving metabolic health, because their recruitment and activation increase energy expenditure and ameliorate obesity-associated metabolic dysfunction^4,6,8,15,51^. In adults, however, beige adipocyte activity is typically transient and heavily dependent on sympathetic stimulation, which limits the durability and translational potential of beige fat-based strategies^5,7–9^. By contrast, during the early postnatal period beige adipocytes arise in murine subcutaneous fat even in the absence of cold exposure, at a time when sympathetic innervation is not yet fully mature^20–26^. Previous studies has shown that beige adipocytes generated in this peri-weaning window can subsequently regress into a dormant state but retain the plasticity to be reactivated, thereby contributing to long-term thermogenic potential and improved metabolic outcomes in adulthood^20,24,27–29^. Nevertheless, it has remained unclear how developmental signals gate this spontaneous beiging and shape the later thermogenic capacity of subcutaneous fat. In this context, our current study identifies Wnt/β-catenin signaling as a developmental brake on beige adipocyte formation. Suppression of this pathway during the peri-weaning period selectively enhances adipocyte thermogenic activation in subcutaneous depots and imprints a higher thermogenic set point that persists into adult life. These findings position developmental modulation of Wnt/β-catenin signaling as a mechanism that links early-life beige adipogenesis to long-term thermogenic capacity.

The canonical Wnt pathway is classically recognized as a negative regulator of adipogenesis^52–54^, and our findings extend this view by showing that Wnt/β-catenin also constrains adipocyte thermogenic potential. In adipocyte-specific β-catenin loss-of-function models, lowering Wnt/β-catenin activity enhances beige-associated oxidative metabolism in iWAT while maintaining adipocyte differentiation markers and systemic adipokine profiles. At the level of lipid metabolism, Wnt/β-catenin inhibition preferentially augments triglyceride hydrolysis and fatty acid oxidative pathways without overtly reducing de novo lipogenesis, indicating a bias in lipid handling towards fuel mobilization and oxidation to support heat production, rather than thermogenesis arising as a secondary consequence of global defects in lipid synthesis or adipose storage described in other thermogenic models^55–57^. This pattern is compatible with a prior report showing that the transcriptional regulator BCL6 is required for the establishment of a beige adipocyte program around the peri-weaning period, partly by supporting fatty acid oxidation and mitochondrial uncoupling^24^, and that BCL6 can antagonize Wnt signaling^58–60^, pointing to potential cross-regulatory interactions between these pathways. In line with this, previous studies have reported that deletion of *Ctnnb1* in adipocytes alleviates long-term high-fat diet-induced metabolic dysfunction, with reduced adipose tissue inflammation and improved insulin sensitivity, features that resemble the systemic benefits typically attributed to increased beige fat activity^61^. Although most mechanistic insights derive from mouse models, human genetic and expression data also support an association between aberrant Wnt activation and adverse metabolic remodeling. For example, gain-of-function alterations in RSPO1-LGR4 pathway in obese individuals have been associated with enhanced Wnt/β-catenin activity and impaired adipose tissue plasticity^62^. Consistent with a conserved role in humans, we observed that pharmacological inhibition of Wnt/β-catenin signaling in primary human subcutaneous adipocytes similarly activates a WNT5A-Ca^2+^-AMPK-related thermogenic response without overt impairment of adipogenic markers. Together with our developmental data, these observations highlight Wnt/β-catenin signaling as a key node in adipocyte plasticity that restricts beige adipocyte thermogenic programming by modulating lipid partitioning and thereby impacts systemic metabolic homeostasis.

Cold exposure is known to recruit beiging in adult iWAT, yet this response is not spatially uniform, and the determinants of such subregional heterogeneity remain unclear^63,64^. We observed a spatial gradient of Wnt/β-catenin activity across iWAT, with higher *Axin2* expression in the posterior region, where beiging was comparatively blunted. Interestingly, β-catenin deletion produced a more pronounced thermogenic shift in this posterior compartment, consistent with greater responsiveness when the canonical Wnt pathway is relieved. These findings raise the possibility that local Wnt/β-catenin activity sets a tissue-intrinsic permissiveness for thermogenic recruitment, potentially interacting with cold-driven extrinsic inputs. Further investigation will require subregion-resolved functional analyses and targeted genetic manipulation.

AMPK is a key metabolic regulator in adipocytes, promoting catabolic and oxidative pathways that support thermogenic responses^12^. Ca^2+^-dependent kinases such as CaMKK2 act upstream of AMPK, and Ca^2+^ flux has been implicated in the activation of beige adipocytes^65–68^. Although the non-canonical Wnt signaling via Wnt/Ca^2+^ has been studied in adipose tissue, work has focused largely on depot-specific traits and inflammatory regulation^69–71^, so its direct contribution to beige thermogenesis has remained uncertain. Wnt5a, a prototypical non-canonical ligand, is a well-established trigger of Wnt/Ca^2+^ signaling and has been linked to metabolic regulation in other contexts^72,73^, but its role in beiging had not been directly addressed. By combining β-catenin loss-of-function with pharmacological manipulation of Wnt5a/Ca^2+^ signaling, we position a Wnt5a-responsive Ca^2^⁺-AMPK-PPAR module downstream of reduced Wnt/β-catenin activity. In peri-weaning iWAT, β-catenin inhibition is accompanied by increased Wnt5a and Fzd5 expression, enhanced phosphorylation of CaMKK2 and AMPK, and coordinated induction of lipolytic and fatty acid oxidative genes. Functionally, interfering with Wnt5a/Ca^2^⁺ signaling attenuates the thermogenic response to β-catenin inhibition, whereas recombinant Wnt5a alone has little effect in the presence of intact Wnt/β-catenin signaling, indicating that Wnt5a-responsive inputs require a lower β-catenin state. This dependence on Wnt/β-catenin activity is consistent with broader developmental paradigms in which the canonical and Wnt5a-dependent non-canonical branches antagonize each other, shaping a hierarchical balance in pathway outputs^30,74^. In addition, receptor context can determine whether Wnt5a dampens or redirects β-catenin/TCF-dependent transcription^75^, aligning with our finding that exogenous Wnt5a is insufficient to engage AMPK and beiging under the canonical Wnt-active conditions. Thus, our findings support a model in which Wnt/β-catenin signaling exerts dominant repression over a Wnt5a-Ca^2^⁺-AMPK-PPAR circuit that can otherwise promote lipid mobilization and oxidative thermogenic gene expression.

In addition to this signaling hierarchy, our data point to temporal tuning of the Wnt5a-Ca^2^⁺ axis. In peri-weaning iWAT, Wnt5a and Fzd5 expression and Ca^2^⁺-AMPK activity are elevated at a time when canonical Wnt signaling is low and spontaneous beige recruitment is robust. In adult *Adipoq*^Cre^;*Ctnnb1*^flox/flox^ mice, however, Wnt5a and Fzd5 transcript levels are no longer markedly different between genotypes, yet CaMKK2 and AMPK phosphorylation and oxidative-lipolytic programs remain higher in iWAT, suggesting that the non-canonical Wnt induction is most prominent during the developmental beiging phase, although sustained AMPK activation and lipid remodeling are likely to contribute to the maintenance of an increased thermogenic tone later in life. Moreover, the absence of a comparable beiging response in BAT or eWAT indicates that this Wnt5a-Ca^2^⁺-AMPK module operates in a depot-restricted manner. Dissecting the precise temporal hierarchy of Wnt5a-Ca^2^⁺-AMPK axis and distinguishing developmental from purely adult roles will require future studies using inducible, stage-specific genetic approaches. Nonetheless, our current findings establish that, during the peri-weaning period, relief of the canonical Wnt/β-catenin signaling is coupled to the activation of a Ca^2^⁺-AMPK-PPAR module that supports beige adipocyte formation and contributes to a higher thermogenic capacity in subcutaneous fat.

## Methods

### Mice

All animal experiments were conducted in accordance with the guidelines and were approved by the Ethics Committee of the Experimental Animal Ethics Committee of West China Hospital of Stomatology, Sichuan University (WCHSIRB-D-2024-560). All procedures were carried out following the Animal Care Committee of Sichuan University. All mice were on a C57BL/6J genetic background. Wild-type mice were purchased from Chengdu Dossy Experimental Animals. *Adipoq*^Cre^;*Ctnnb1*^flox/flox^ mice were generated by compounding *Adipoq*^Cre^ (Stock No. 028020; Jackson Laboratory) allele with *Ctnnb1*^flox^ allele (Stock No. 004152; Jackson Laboratory). *Ucp1*^Cre^;*Ctnnb1*^flox/flox^ mice were generated by compounding *Ucp1*^Cre^ (Stock No. 024670; Jackson Laboratory) allele with *Ctnnb1*^flox^ allele (Stock No. 004152; Jackson Laboratory). *Adipoq*^Cre^;*Ctnnb1*^dm/flox^ mice were generated by compounding *Adipoq*^Cre^ (Stock No. 028020; Jackson Laboratory) allele with *Ctnnb1*^flox^ allele (Stock No. 004152; Jackson Laboratory), and *Ctnnb1*^dm^ allele (gift from Dr. K. Basler of the University of Zurich). Unless stated, all experiments were performed on male mice aged 3–8 weeks. Mice had free access to food and water and were housed under a 12-hour light–dark cycle with controlled temperature and humidity (approximately 22 °C and 60% humidity). Primer sequences for genotyping are listed in Supplementary table 2.

### Isolation of mouse adipose stromal vascular fractions (SVFs)

All cells were isolated from adult C57BL/6J male mice, unless otherwise specified. For the isolation of adipose SVF cells, inguinal fat pads were carefully dissected and transferred into sterile phosphate-buffered saline (PBS), then rinsed sequentially in PBS containing 5% Penicillin-Streptomycin (PS) (15140122, Gibco), 2% PS, and finally in sterile PBS, each wash performed 2–3 times to remove blood and contaminants. The cleaned tissue was minced into small fragments and transferred into T75 culture flasks. After allowing the tissue to settle and adhere for 5–10 minutes, complete culture medium (a-MEM with 10% fetal bovine serum and 1% PS) was gently added to each flask.

### In vitro pro-adipogenic differentiation of mouse cells

For mouse adipocyte differentiation, isolated adipose SVF cells were seeded into plates at the density of 1 × 10^4^ cells/cm^2^ in complete culture medium and treated with Mouse Adipose-derived Mesenchymal Stem Cells Adipogenic Differentiation Cocktail (MUXMD-90031, OriCell) with 90% cell confluence. Cells were cultured for several days until complete adipogenic differentiation was observed in the entire cell population. For inhibition and activation experiments, once full adipogenic differentiation was confirmed, cells were treated with each of the following molecules for 4 days: MSAB (S6901,Selleck, 10 uM), BAPTA-AM (S7534, Selleck, 5 uM), Box5 (P1216, Selleck, 100 uM), Wnt5a recombinant protein (HY-P704313, MCE, 100 ng/mL) with the exception of Rosiglitazone (S2556, Selleck, 2 uM), which was added on the first day of adipogenic induction in SVFs. In all experiments, vehicle groups were treated with DMSO at a fixed concentration of 0.1% (v/v), which was equal to the concentration present in all drug-treated groups.

### Isolation and pro-adipogenic differentiation of human subcutaneous adipocytes

Human adipose tissue was collected from the abdominal or thigh regions of patients undergoing liposuction at Chengdu Junda Surgical Plastic Hospital. Eligible donors were between 18 and 35 years old and had normal blood pressure and blood glucose levels. Individuals with any infectious or vascular disorders were excluded. All samples were obtained with informed consent from participants. All experimental procedures were conducted under the guidelines of the Ethics Committee of West China Hospital of Stomatology, Sichuan University, and approved by the Ethics Committee (WCHSIRB-D-2025-560). The tissue was washed with PBS to remove blood cells, minced into fine pieces, and digested with Type I collagenase (C0130, Sigma-Aldrich, 1 mg/mL) at 37°C for 30 minutes in a shaking incubator at 150 rpm. After the digestion was stopped, the sample was left to stand for 5 minutes. The floating adipocytes and top lipid layer were removed. The remaining cell suspension was filtered through 500 μm and 40 μm mesh filters to separate and purify the SVFs. Finally, the filtrate was centrifuged at 300 × g for 5 minutes to pellet the cells. Isolated human SVF cells were seeded at a density of 1 × 10^4^ cells/cm². Upon reaching 90% confluence, the culture medium was replaced with Human Adipose-derived Mesenchymal Stem Cells Adipogenic Differentiation Cocktail (HUXMD-90031, OriCell) to induce differentiation. Following complete adipogenic differentiation, the cells were treated with 10 μM MSAB (S6901, Selleck) for 4 days.

### Hematoxylin and eosin staining and cell counting

Adipose tissues were fixed in 4% paraformaldehyde (PFA) and embedded in paraffin. Tissue sections (7 μm) were subjected to hematoxylin and eosin (H&E) staining following standard procedures. HE images from iWAT were used for morphometric analysis. Cell counting was achieved using the ImageJ software.

### Immunofluorescence

Freshly isolated adipose depots were fixed in 4% PFA at 4 °C overnight, followed by embedding and sectioning. For immunostaining, paraffin-embedded tissue sections were deparaffinized in xylene and subsequently rehydrated through graded ethanol solutions to PBS. After incubating the sections in boiling Tris-EDTA buffer (C1034, Solarbio) for 5 minutes, the slides were washed and blocked in PBS containing 10% Bovine Serum Albumin (BSA) (V900933, Sigma-Aldrich) for 30 minutes and then incubated with the primary antibodies against Ucp1 (Abcam, ab23841, 1:300), Perilipin (Invitrogen, MA5-27861, 1:500) at 4 °C overnight. Slides were then incubated with secondary antibodies (Alexa Fluor 488-anti mouse, A21202; Alexa Fluor 647-anti rabbit, A31573; Invitrogen, both 1:500) at room temperature for 30 min, followed by staining with 4’,6-diamidino-2-phenylindole (DAPI, D8200, Solarbio). For immunostaining on cell culture, cells were fixed with 4% PFA for 15 min at room temperature and then stained with Nile Red (C2051, Beyotime) and DAPI for 10 min at ambient temperature respectively. All images of samples were captured with Olympus confocal Microscope FV2000 and analyzed by using the ImageJ software.

### RNA preparation and quantitative RT-qPCR

Total RNA was extracted from tissues or cells using FastPure Cell/Tissue Total RNA Isolation Kit V2 (RC112-01, Vazyme) according to the manufacturer’s instructions. The concentration and quality of RNA were determined by NanoDrop 2000 (Thermo). Reverse transcription was performed with the HiScript III RT SuperMix for qPCR (+gDNA wiper) (R32301, Vazyme). RT-qPCR was performed using Tap Pro Universal SYBR qPCR Master Mix (Q712, Vazyme) on a QuantStudio 6 Flex detection system. Values of each gene were normalized to reference genes 36B4, using the 2^−ΔΔ^Ct method. Primer sequences are listed in Supplementary table 3.

### Western blotting

Proteins were extracted from adipose tissue or cultured adipocytes using Adipose Tissue Protein Extraction Kit (EX1130, Solarbio) according to manufacturer’s instructions. Lysis buffer also contained Protease and phosphatase inhibitor cocktail (P1045, Beyotime). The protein concentrations were measured using BCA protein assay kits (KGP902, KeyGEN BioTECH). Subsequently, proteins were separated by SDS-PAGE and transferred onto PVDF membranes (ISEQ00010, Millipore). Membranes were blocked in 5% skim milk or 10% BSA for 1 h, followed by overnight incubation with primary antibodies. Antibodies were purchased and diluted as follows: anti-UCP1 (1:1000); anti-OXPHOS (Abcam, ab110413, 1:500); anti-β-catenin (Santa Cruz, sc-59737, 1:1000); anti-Adiponectin (Abcam, ab22554, 1:1000); anti-β-actin (Proteintech, 60008-1-Ig, 1:5000); anti-Wnt5a (Zen, 619919, 1:1000); anti-pCaMKII (Thr286) (Abcam, ab32678, 1:1000); anti-CaMKII (Abcam, ab52476, 1:1000); anti-pCaMKK2 (Ser511) (CST, 12818, 1:1000); anti-CaMKK2 (Proteintech, 11549-1-AP, 1:1000); anti-pAMPKα (Thr172) (CST, 2535, 1:1000); anti-AMPKα (Proteintech, 10929-2-AP); anti-PPARα (Abcam, ab227074, 1:1000); anti-ATGL (CST, 2138, 1:1000); anti-CPT1 (Santa Cruz, sc-393070, 1:1000); anti-Sirt1 (Zen, 617857, 1:1000);anti-Pgc1α (BBI, D262041, 1:1000); anti-PPARγ (Abcam, ab272718, 1:1000); anti-Axin2 (Abcam, ab109307, 1:1000); anti-LEF1 (Abcam, ab85052, 1:1000); anti-Active beta-catenin (Ser33/37/Thr41) (CST, 8814, 1:1000). Samples were then incubated with the corresponding horseradish peroxidase-conjugated secondary antibodies (Anti-Rabbit IgG H&L, 511203; Anti-Mouse IgG H&L, 511202, Zen, both 1:5000) for 1 hour. Blots were visualized using an ECL detection kit (180–5001, Biotanon), and quantitative analysis was performed using ImageJ software.

### Assessment of mitochondrial respiration

Oxygen consumption rates of iWAT and cultured differentiated adipocytes were measured with Seahorse XFe24 analyzer (Agilent Technologies). For adipose tissue, isolated iWAT was placed in wash medium (Seahorse XF DMEM supplemented with 25 mM glucose and 25 mM HEPES) for preconditioning. Tissue cores (∼4 mg) were then obtained from the middle region using a 2 mm biopsy punch and placed into wells of an XF24 Islet Capture Microplate (Agilent Technologies, 101122-100), where the capture screen was gently secured to ensure the tissue in place. Each well was filled with 450 μl of assay medium (Seahorse XF DMEM supplemented with 25 mM glucose) and incubated at 37 °C in non-CO_2_ environment for 1.5 hours. Samples were subjected to a mitochondrial stress test by adding oligomycin (GC16533, GlpBio, 12 μM), FCCP (C2920, Sigma-Aldrich, 8 μM), rotenone (R8875, Sigma-Aldrich, 12 uM) and antimycin A (GC49360, GlpBio, 12 μM) according to the manufacturer’s instructions. For cells, SVFs were seeded in an XF24 cell culture microplate (Agilent Technologies, 102342–100) at a density of 20,000 cells per well. After 7 days of adipogenic induction and 4 days of molecule treatment, 500 μl XF assay medium containing 1 mM pyruvate, 2 mM glutamine, and 10 mM glucose was added to each well. Mitochondrial stress test was performed on cells by adding oligomycin (5 μM), FCCP (2 μM), and rotenone and antimycin A (5 μM).

### Serum adiponectin and biochemical measurements

Blood was collected by cardiac puncture upon euthanasia. After coagulation at room temperature for 2 h and centrifugation at 2,000 × g for 15 min at 4 °C, separated serum was transferred to a new tube and stored at -80 °C until use. Serum adiponectin, triglyceride, total cholesterol and free fatty acid levels were measured using commercial enzyme-linked immunosorbent assay kits according to the manufacturer’s instructions (Jonln Bio, Shanghai, China).

### Metabolic cages

Whole-body energy metabolism was evaluated using a comprehensive laboratory animal monitoring system home cage (CLAMS-HC) (Columbus Instruments) following manufacturer’s protocol on 8-week-old littermate *Adipoq*^Cre^;*Ctnnb1*^flox/flox^ and control mice. Each mouse was housed in individual chambers for 6 days. Each parameter was measured every 5 min. EE and RER were calculated based on oxygen consumption (O_2_) and carbon dioxide production (CO_2_). Food intake was measured via weight sensors.

### Bulk RNA-Seq

For RNA sequencing of tissue samples, freshly isolated iWAT was immediately snap-frozen in liquid nitrogen for 30 minutes. For cellular sequencing, SVF from 4-week-old and 8-week-old wild-type mice were differentiated into mature adipocytes in vitro, then lysed with TRIzol regnant (9109, Takara, Japan) and snap-frozen in liquid nitrogen for 30 minutes. All samples were frozen at -80 °C before RNA extraction and analysis. The following steps were conducted by Majorbio Bio-Pharm Technology Co., Ltd. (Shanghai, China). Total RNA was isolated from the samples, followed by genomic DNA removal using DNase I (Takara, Japan). RNA-seq libraries were generated from 1 μg of total RNA using the TruSeq™ RNA Sample Preparation Kit (Illumina, San Diego, CA, USA) according to the manufacturer’s instructions. In brief, polyadenylated mRNA was enriched, fragmented, and reverse-transcribed into double-stranded cDNA using the SuperScript Double-Stranded cDNA Synthesis Kit (Invitrogen, CA) with random hexamer primers (Illumina). The resulting cDNA fragments underwent end-repair, phosphorylation, and 3′-end adenylation. After fragment size selection (∼300 bp), sequencing adapters were ligated, and the libraries were subjected to paired-end sequencing (2 × 150 bp) on an Illumina HiSeq X Ten or NovaSeq 6000 platform. All bioinformatics analyses were conducted on the Majorbio Cloud Platform (www.majorbio.com).

### Statistical analysis

Data collection and analysis were not performed blind to the conditions of the experiments. Data were shown as mean ± SEM and were analyzed with GraphPad (v.10.0.3). Statistical differences between two-groups were calculated by two-tailed unpaired Student’s *t*-test. Multiple group comparisons were conducted using one-way ANOVA followed by Tukey’s test. For Seahorse measurement and energy metabolism analysis, two-way repeated measures ANOVA was performed with Bonferroni’s correction. For all experiments *P* < 0.05 was considered significant.

## Data availability

The RNA-seq dataset has been deposited in the GEO database (GSE315581, GSE315582, GSE315583). Additional data supporting the plots are available in the Supplementary Information or Source data files.

## Acknowledgements

We thank members of the Chen Lab for sharing reagents. We thank Dr. Konrad Basler of the University of Zurich for his kind gift of *Ctnnb1*^dm^ mice. This work was supported by National Key Research and Development Program of China (2022YFA1104400), Sichuan Science and Technology Program (2024YFFK0068, 2026YFHZ0046), and Research and Develop Program, West China Hospital of Stomatology Sichuan University (RD-02-202517).

## Author contributions

J.X., B.D., Y.C., W.T., T.C., and Z.L. conceived and designed the study. G.X. and T.C. performed experiments, data analysis and prepared the manuscript. Q.Z., B.Z. and J.X. assisted in performing experiments. W.T. and T.C. provided funding and, together with Z.L., supervised the study. All authors provided intellectual input and assisted with the preparation of the manuscript.

## Additional information

Supplementary Figures.

Table 1. Reagent/chemical information.

Table 2. Primer sequences used for genotyping.

Table 3. Primer sequences used for quantitative RT-qPCR.

